# Computational design of intrinsically disordered protein regions by matching bulk molecular properties

**DOI:** 10.1101/2023.04.28.538739

**Authors:** Bob Strome, Khaled Elemam, Iva Pritisanac, Julie D. Forman-Kay, Alan M. Moses

**Affiliations:** Departments of Cell & Systems Biology, University of Toronto; Departments of Biochemistry, University of Toronto; Departments of Hospital for Sick Children Research Institute, Toronto

## Abstract

Algorithms for computational protein design usually begin with a 3D structure and design an amino acid sequence that will fold into that structure. Since intrinsically disordered protein regions lack stable 3D structures, strategies for their design have been limited. Here we describe and validate a general computational strategy for design of intrinsically disordered regions (IDRs). Our algorithm designs synthetic IDRs by minimizing the distance of an initially random amino acid sequence to a known IDR in a high-dimensional space of molecular properties, such as repeat content, charge and hydrophobicity. We tested a handful of IDRs designed to target proteins to mitochondria and heat-induced condensates, and several showed the expected patterns in cells.

**One sentence summary:** We design synthetic mitochondrial targeting signals and condensate forming intrinsically disordered regions

## Introduction

Recently, computational protein design algorithms have achieved practically useful performance for folded protein regions [1–4] and several groups have been exploring computational approaches to design intrinsically disordered regions (IDRs) [5–7]. In folded protein design, proteins are usually designed to match a pre-specified tertiary structure[8]: because of the strong connection between folded protein structure and biological function, if the structure is achieved, the designed protein can be expected to perform its biological function. On the other hand, because IDRs often function through dynamic ensembles and do not take on stable tertiary structures[9,10], a general strategy for computational protein design for IDRs has been less clear. IDR design efforts have therefore been customized for specific biological functions, such as linkers[5], formation of membrane-less organelles[7,11–14], targeting signals[15,16] or interaction domains[17].

Although they do not fold into stable structures, there is increasing evidence that bulk molecular properties such as charge, hydrophobicity, motif and repeat content allow intrinsically disordered regions (IDRs) to perform a variety of key functions in the cell in their natively disordered conformations[18–22]. For example, we showed that, when considered in combination, bulk molecular properties are sufficient to predict many biological functions of IDRs[21]. Furthermore, we were able to predictably modulate in vivo function of IDRs by adding and removing charge and hydrophobicity[21,23]. If bulk molecular properties determine IDR function, we hypothesize that bulk molecular properties could be used as the analog of tertiary structure for general computational IDR design (Figure S1).

Here we report our first attempts at computational IDR design using an algorithm that designs peptides to match the bulk molecular properties of extant protein regions. First, we aimed to design a mitochondrial localization signal by matching the bulk properties of the N-terminus of Cox15, previously identified as a mitochondrial pre-sequence[24] and necessary for mitochondrial localization[21]. Remarkably, when we tested 3 of these designed IDRs in cells, we found that two of them showed co-localization with the native Cox15. Second, we tried to design IDRs that support heat dependent protein condensation[25] by matching the properties of IDRs from Ded1. We found that our C-terminal design was sufficient for targeting to Ded1 droplets, while the N-terminal design could not. Finally, we designed an IDR that had molecular properties found in certain condensate forming IDRs. We found that it showed constitutive condensates that were distinct from the heat-dependent Ded1 condensates.

Taken together these results strongly support the hypothesis that IDR function is determined by the bulk molecular properties, and that it is relatively straightforward to design amino acid sequences that can encode these bulk properties. Our results suggest that design of IDRs with specific biological functions may be a much easier problem than design of folded proteins.

## Results

### Design of a mitochondrial localization signal

We first attempted to design the intrinsically disordered 45 residue mitochondrial localization signal (also known as a presequence or mitochondrial targeting signal) from the *S. cerevisiae* protein Cox15 (we refer to as the design “target”, Figure 1a). Although they can form amphipathic helices upon binding to their receptors[26], mitochondrial localization signals are increasingly appreciated to bind in multiple conformations in dynamic equilibrium [27], typical of many intrinsically disordered interactions[28]. We collected 94 molecular properties that could be computed directly from the primary amino acid sequence (Table S1) and computed these for the Cox15 N-terminal IDR. We refer to this as the “target molecular property vector.” Next, starting with a random peptide of 45 amino acids, at each iteration, we selected the single substitution that led to the greatest reduction in the Euclidean distance between the scaled molecular property vectors (See Methods) of the designed peptide (design) and the target molecular property vector. This greedy optimization strategy resulted in designs that showed little primary sequence similarity (Figure 1b, see Methods) to the Cox15 N-terminus, albeit more than the random initial peptides (e.g., for n=10 designs, average pairwise % identity is 18.6 for the designs and 11.2 for the random initial peptides, P<0.05 paired T-test) but much less than orthologous Cox15 N-termini from closely related fungi (*sensu stricto* Saccharomyces and more diverged homologs, with average pairwise % identity of 78.5 and 28.9 for n=3 and n=15, respectively, T-test, P<0.005 for both, see Methods).

**Figure 1.**
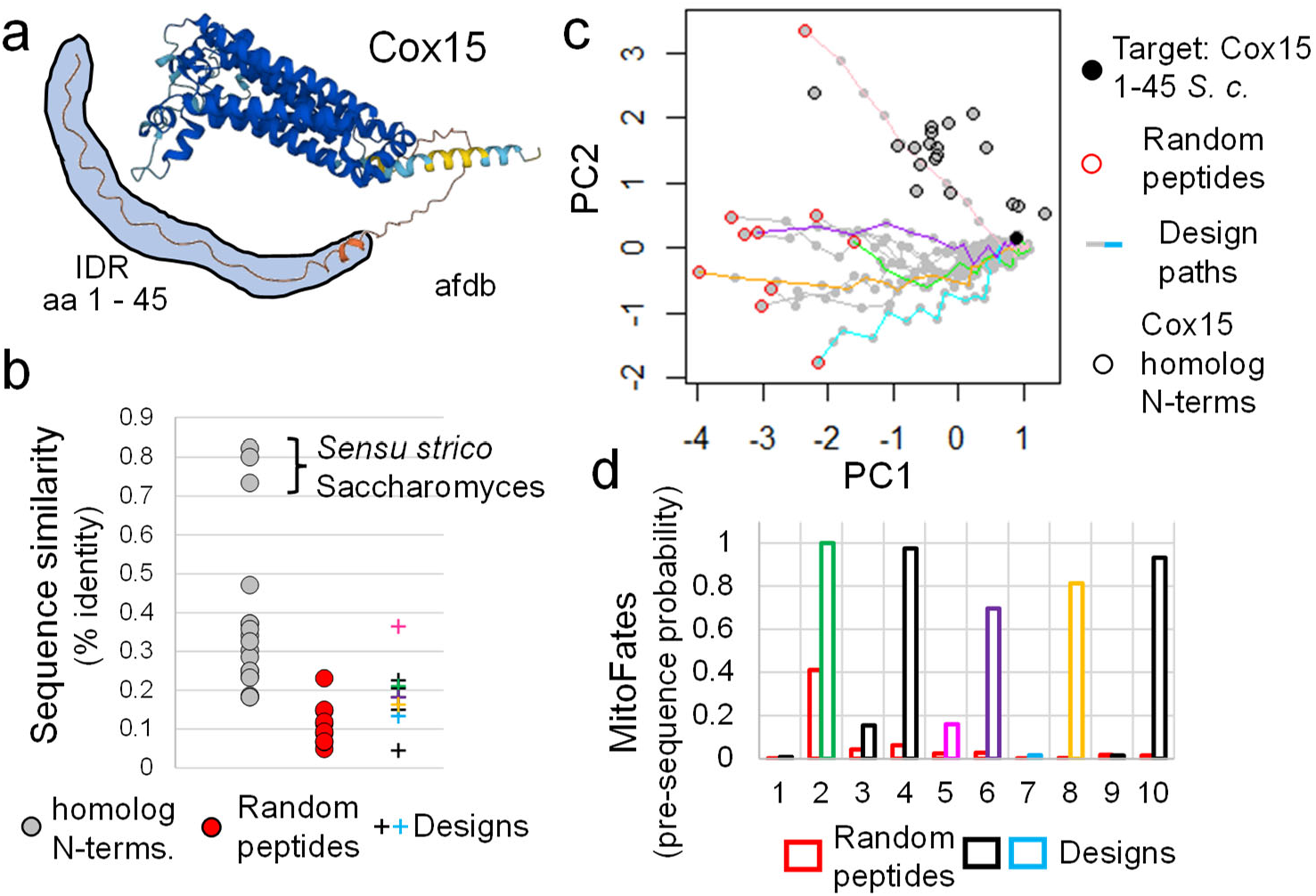
Design of Cox15 N-terminal IDRs a. *S. cerevisiae* Cox15 structure from AlphaDoldDB [30] (afdb), showing the extended N-terminus (highlighted with light blue). b. pairwise sequence identity of homologous Cox15 N-termini (grey symbols), random initial peptides (red symbols) and designs (coloured crosses) to the design target (*S. cerevisiae* Cox15 N-terminus). c. Ten example design series (points connected by colored or grey lines), starting from the random initial peptides (red outlines) towards the target (filled symbol) are shown. Peptides (grey points) are represented by their loadings on the first and second principal components (PC1, PC2, respectively) of the molecular properties space. For reference, N-termini of Cox15 homologs are shown (black outlines). d. examples of MitoFates predictions for random initial random peptides (red bars) and the designs (black and coloured bars) after greedy optimization with Cox15 as the target. Y-axis represents the probability of possessing a pre-sequence computed by MitoFates.

To test whether the greedy algorithm could reach sequences with molecular properties “close enough” to the target, we compared them to the naturally occurring Cox15 N-termini from closely related fungi (unfilled symbols in Figure 1c) that we expect to have the same function as the target because Cox15 mitochondrial localization is conserved across eukaryotes. To compare the molecular properties, we plotted “design paths” starting from the random peptides (red symbols in figure 1c) and approaching the target (filled symbol in figure 1c) along with the homologs in a reduced dimensional space defined by the first two principal components of the 94 bulk molecular properties. The designed sequences were closer to the target in this space than homologs from related fungi (Figure 1c), indicating that the greedy algorithm had identified very similar sequences in the molecular property space. This is consistent with the idea that large numbers of peptide sequences can encode the molecular properties of the Cox15 N-terminus. We reasoned that if the 94 molecular properties we defined included the sufficient properties for a mitochondrial pre-sequence, the level of similarity in molecular property space achieved by the greedy algorithm would likely be sufficient for function.

To test the function of the designed sequences, we replaced the N-terminus of Cox15 with the designs (or random initial peptides) and submitted them to MitoFates[29], a state-of-the-art sequence-based predictor of mitochondrial localization signals, which assigns an input protein sequence a probability of possessing a pre-sequence. We found that many of the designed sequences (31 of 50) and only one of the random initial peptides were predicted to possess pre-sequences (P-value < 10^−6^, Fisher’s exact test, Figure 1d shows 10 examples of which 5 of the designed IDRs are predicted to possess presequences) suggesting that a majority of the designed peptides are likely to function as mitochondrial localization signals. The sequences of these IDR designs all display similar enrichments of hydrophobic and positively charged residues, known hallmarks of mitochondrial localization signals. Three designed peptides are shown in Figure 2a, two of which were predicted to be mitochondrial pre-sequences by MitoFates (P1 and P2), while the other was not (N1).

**Figure 2.**
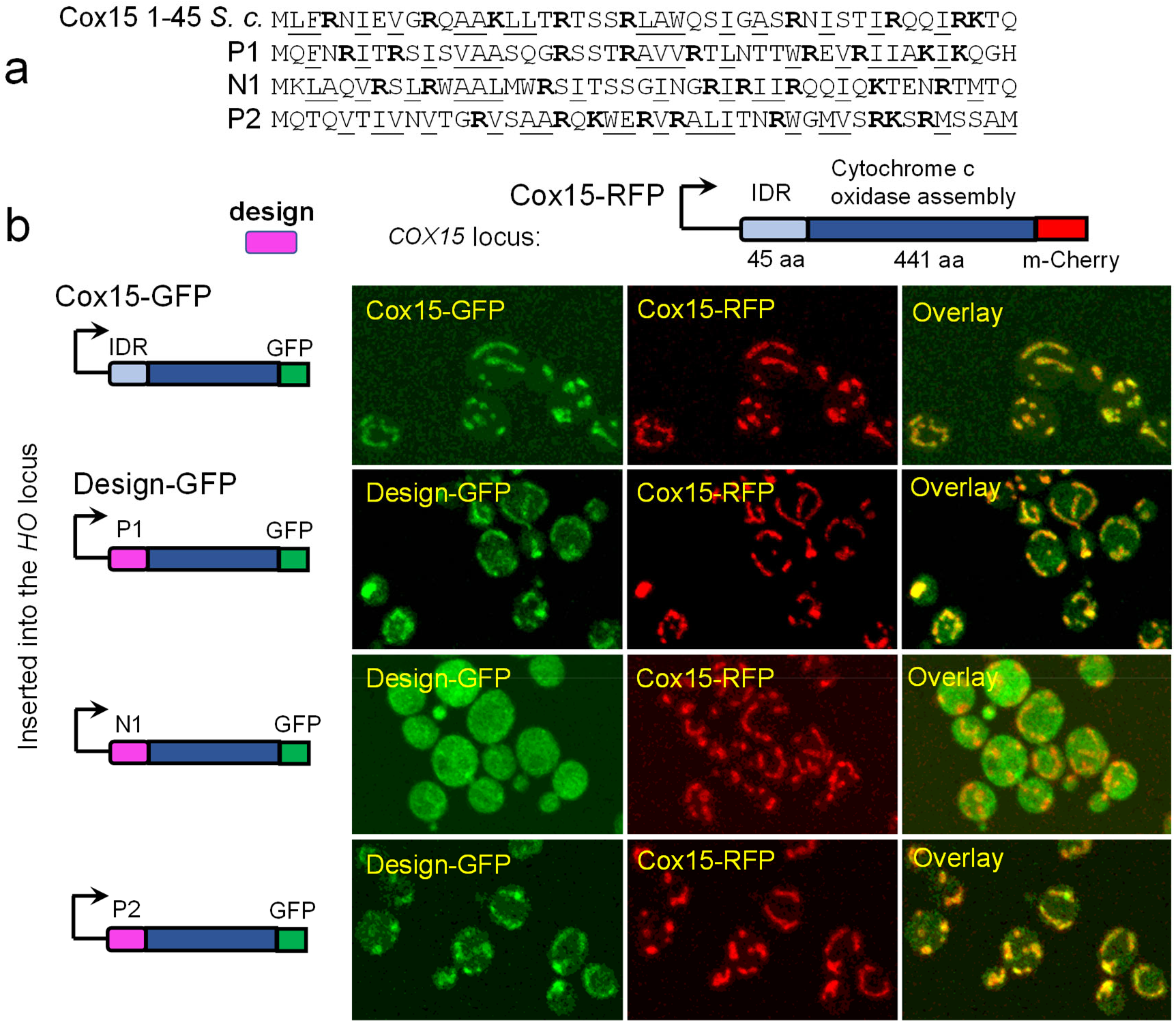
Expression of synthetic mitochondrial localization signals a. Sequence of the design target, wild-type Cox15 N-terminus, and three computational designs generated by our greedy algorithm that matches the target molecular properties. Basic (bolded) and hydrophobic (underlined) residues are indicated. b. Confocal fluorescence microscopy images of cells expressing of GFP-tagged Cox15 with wildtype and designed IDRs (P1, N1, and P2, see text for details) at the *HO* locus and RFP-tagged Cox15 at the endogenous locus. Each row represents a different strain where a schematic of the fluorescent tagged protein expressed is shown on the left.

Next, we constructed yeast strains that expressed full length GFP-tagged Cox15 where we replaced the N-terminus of with three of our designed IDRs (Figure 2a) at a second locus (the *HO* locus). In these strains, we tagged the endogenous copy of the Cox15 tagged with RFP that allowed us to visualize the target localization pattern (See Methods). Remarkably, the two designed peptides predicted by MitoFates showed clear co-localization (Figure 2b, see Methods) with the target, Cox15, indicating that the computational design of IDRs was successful. Interestingly, the designed peptide predicted not to be a targeting signal by MitoFates showed a cytoplasmic pattern. This suggests that there are other sequence-encoded features that were not present in our molecular property space that are recognized by MitoFates. However, these missing sequence-encoded features must be relatively easy to obtain by chance, because we used the same algorithm to design all three of these synthetic IDRs.

### Design of a heat-induced, condensate-forming IDR

Encouraged by the success at designing the mitochondrial localization signal from Cox15, we next focused on a more challenging IDR design problem: the heat-induced condensation of *S. cerevisiae* Ded1[25]. Ded1 contains folded RNA binding domains and N- and C-terminal IDRs (Figure 3a) and we first needed to establish if the IDRs were sufficient for condensation in vivo. To test this, we expressed GFP-tagged truncated versions of Ded1 (either both IDRs only, Figure 3a, top row) or the N-(Figure 3b, second row) and C-terminal IDRs alone at a second locus in cells where the endogenous copy was tagged with RFP. For the C-terminal alone, we were not able to observe convincing GFP-expression, but a construct containing the two IDRs co-localized to the endogenous Ded1 droplets after heat shock, while the construct containing only the N-terminal IDR could not. We noted in both these constructs aberrant nuclear localization (indicated with white arrows in top row) suggesting that we have missed an important nuclear export signal in the construct, or that they are too small to be retained in the cytoplasm by size. Regardless, these experiments confirm that the two Ded1 IDRs together are sufficient to target GFP to the heat-shock dependent Ded1 droplets.

**Figure 3.**
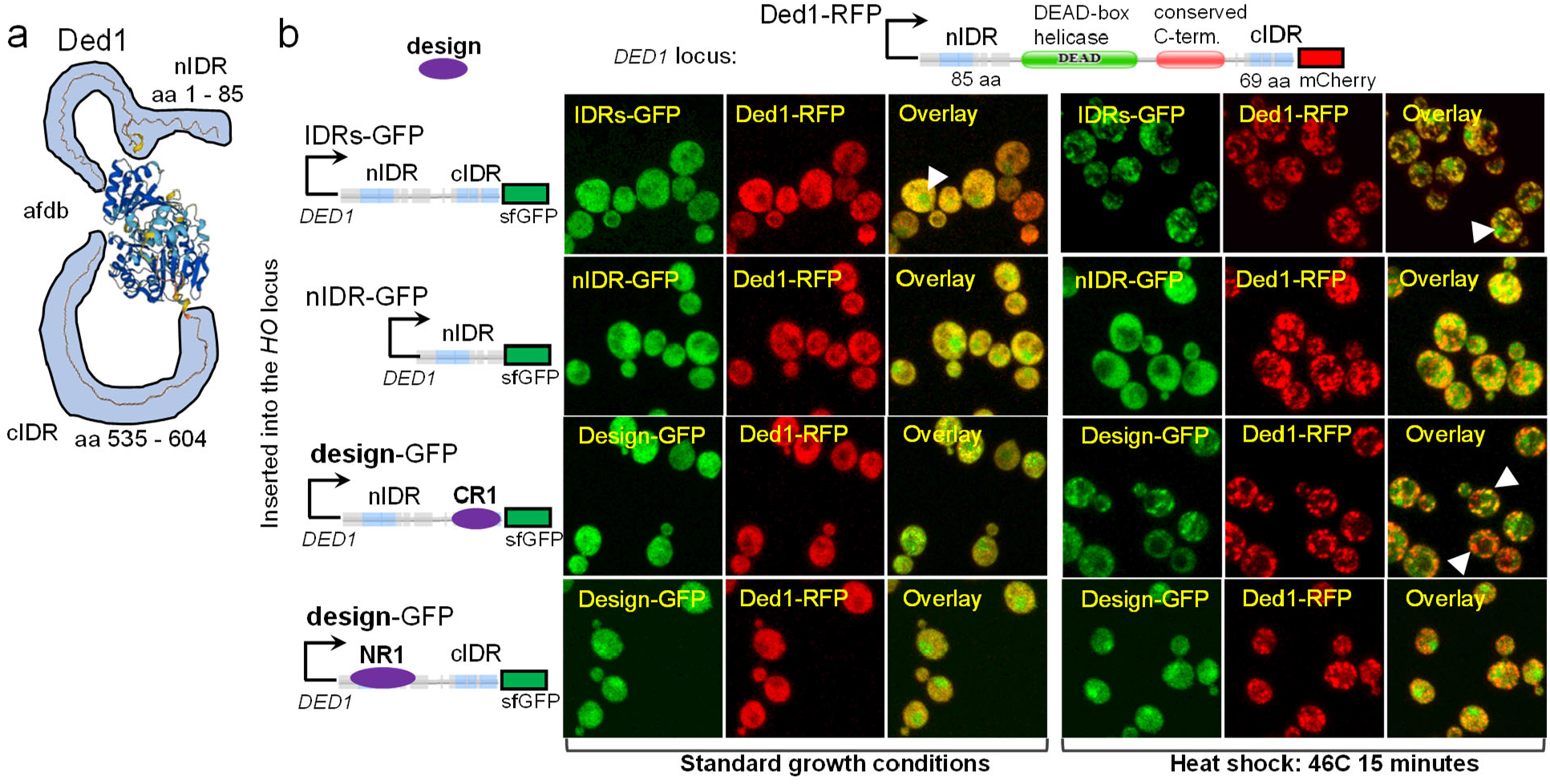
Design of a heat-dependent condensate forming IDR a. the predicted structure of Ded1 from AlphaFold DB (afdb) shows the N- and C-terminal IDRs (highlighted in light blue). b. Confocal fluorescence microscopy images of cells expressing of GFP-tagged IDR constructs (see text for details) at the *HO* locus and RFP-tagged Ded1 at the endogenous locus. Left 3 panels represent standard growth conditions, while right 3 columns represent 15 minutes heat shock at 46°C. Each row represents a different strain, with a schematic of the fluorescently tagged protein expressed is shown on the left. sfGFP: superfolding GFP.

We used the same greedy algorithm to design synthetic peptides that matched the bulk molecular properties of each of the C- and N-terminal IDRs (CR1 and NR1 respectively). Remarkably, we found that the C-terminal design showed at least some co-localization with the Ded1 droplets (Figure 3b, third row, white arrow) although the localization was not as reliable as the wild-type IDRs, as there were cells where the droplets were mostly red only (Figure 3b, third row, white arrow). On the other hand, the N-terminal design (NR1) showed no evidence of co-localization with the Ded1 droplets. Nevertheless, these results indicate that simply by matching the molecular properties of the Ded1 C-terminal IDR, we were able to design an IDR that can specifically target to Ded1 droplets under heat shock.

### Design of a constitutive condensate forming IDR

Although consistent with previous reports that the Ded1 N-terminus is not sufficient for condensation in vitro[25], our observation above that the N-terminal IDR alone was not sufficient to localize to condensates is surprising given that it contains sequence features that are strongly associated with condensation. For example, PScore (a predictor of phase-separation based on Pi-Pi contacts[31]) assigns a high propensity for phase separation to both the N- and C-terminal IDRs (blue trace, Figure 4a). We hypothesized that there may be some additional molecular properties in the Ded1 N-terminus that prevent its condensation[25].

**Figure 4.**
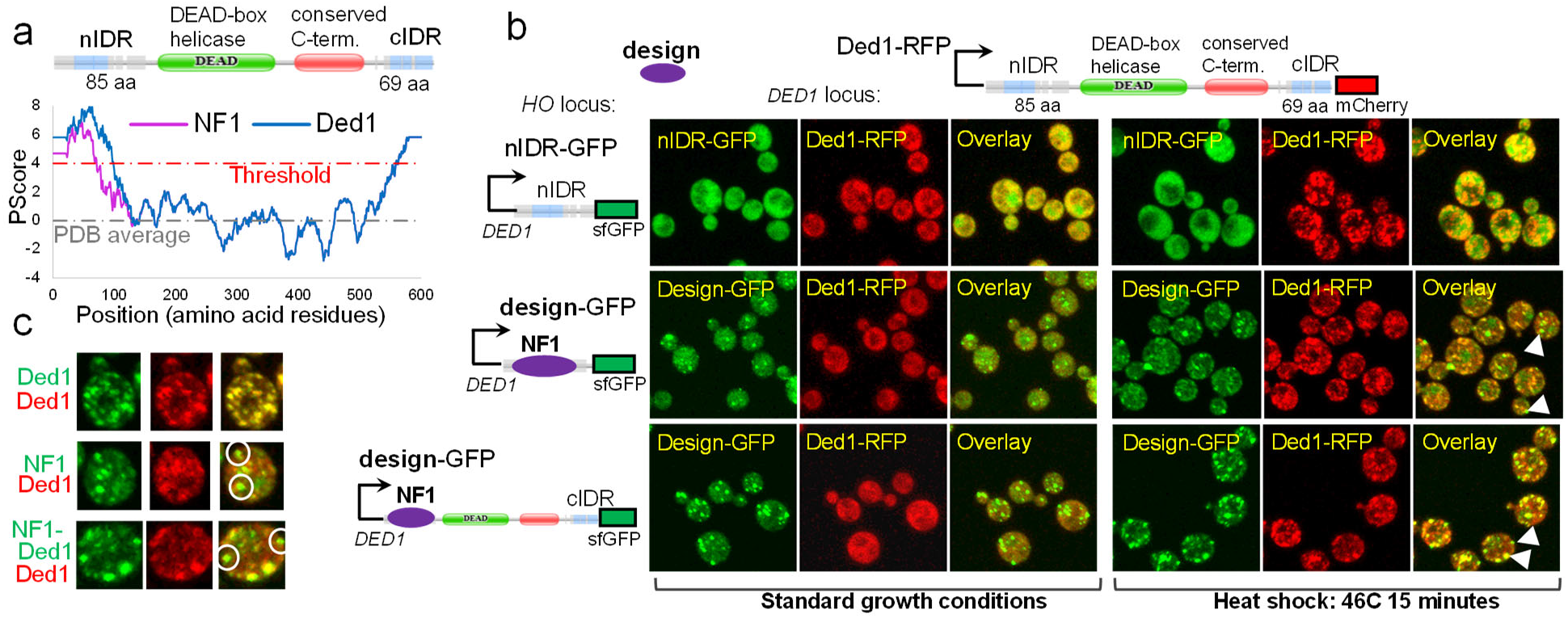
Design of a constitutive condensate forming IDR a. The PScore crosses the threshold (red dashed line) for phase separation prediction for both the N- and C-terminal IDRs of Ded1 (blue trace) as well as when the N-terminus is replaced by a designed peptide (NF1) to match 10 of the molecular properties of Ded1 (Pink trace). b. Confocal fluorescence microscopy images of cells expressing of GFP-tagged IDR constructs (see text for details) at the *HO* locus and RFP-tagged Ded1 at the endogenous locus. Left 3 panels represent standard growth conditions, while right 3 columns represent 15 minutes heat shock at 46 degrees. Each row represents a different strain where a schematic of the fluorescent tagged protein expressed is shown on the left. sfGFP: superfolding GFP. c. Zoom in view of example cells. Top row shows a single cell showing overlap of Ded1-GFP and Ded1-RFP condensates. Second row shows GFP-tagged designed IDR (NF1) condensates that do not overlap with Ded1-RFP condensates (white circles). Third row shows condensates of the GFP-tagged designed IDR replacing the N-terminus of the full length Ded1 (NF1-Ded1) that do not overlap with the Ded1-RFP condensates (white circles). Images are taken after 15 minutes at 46°C.

We reasoned that a synthetic peptide designed only to match the condensation associated properties of the Ded1 N-terminus, might show condensation in cells. To test this, we identified 10 molecular properties that were associated (see Methods) with a collection of condensate localizing IDRs (including G-repeats, RG-repeats, R/K-ratio, fraction of charged residues and a measure of clustered charges, Table S2). We then designed peptides to match the quantitative values of the N-terminus of Ded1 for only these 10 “condensate-associated” molecular properties. As expected, when we replaced the N-terminus of Ded1 with one of these designed peptides (NF1), the PScore for this region remained well above the threshold (pink trace, Figure 4a).

Once again, we tagged NF1 with GFP and expressed it in Ded1-RFP cells. In contrast to the naturally occurring Ded1 N-terminal IDR which was cytoplasmic in standard growth conditions and heat shock (Figure 4b, top row), we found that this designed peptide formed condensates in both standard growth conditions and under heat shock (Figure 4b, second row). These condensates did not fully co-localize with the endogenous heat-dependent Ded1 condensates (Figure 4b, second row, white arrows and Figure 4c, NF1, white circles). We next tested the localization of full length Ded1 when the N-terminus was replaced by NF1, the designed IDR (NF1-Ded1). We found that NF1 was sufficient to localize Ded1 to condensates under standard growth conditions, and that during heat shock, this construct localized to both endogenous Ded1 condensates and additional condensates that do not contain endogenous Ded1 (Figure 4b, third row, white arrows and Figure 4c, NF1-Ded1, white circles). These experiments show that design of IDRs capable of condensation in cells can be achieved with a simple algorithm to match the molecular properties of IDRs.

## Discussion

We used a greedy algorithm to design peptides that simply match the bulk molecular properties of intrinsically disordered regions and found that several of them could support the expected function in cells. The success of a simple greedy algorithm suggests that the IDR design problem is much easier than that of folded proteins, and that there are likely large numbers of functional solutions for IDRs. The relatively loose constraint on IDR sequences is also consistent with the rapid rate of sequence divergence and relative intolerance to point mutations observed for these protein regions[32].

Taken together, our attempts at automatic computational IDR design show that bulk molecular properties of the primary amino acid sequence can be sufficient for function, at least for the subcellular localization functions here (localization to mitochondria, Ded1 droplets and spontaneous formation of droplets). These results are consistent with observations by us and others that these properties are conserved during evolution[23,33,34]. The basis of specificity in biomolecular condensate formation is currently an area of intense research interest in biophysics and cell biology. Our success at designing synthetic condensate forming IDR that forms condensates (NF1) distinct from the Ded1 condensates (Figure 3c) supports the idea that the sequences of IDRs are sufficient to specify distinct condensates in cells[35,36].

Our study leaves many questions open about IDR design. First, not all of the synthetic IDRs we designed showed the expected subcellular localization patterns, and in some cases the patterns they drove were not as clear as the native IDRs. For mitochondrial localization signals, MitoFates, a state-of-the-art bioinformatics predictor, appears to be able to recognize the difference between our functional and non-functional designs. This strongly suggests that there are more complex or additional bulk molecular properties that were not included in our analysis, Experimental and computational methods to systematically discover bulk molecular properties that are important for IDR function could identify these missing features and directly improve IDR design. Here we used a set of 94 properties that we curated from the literature, but recent language-modelling and other deep learning-based approaches can project proteins into feature spaces an order of magnitude larger[37,38]. Designing IDRs in these feature spaces is a promising direction for future research. For the designed Ded1 C-terminal IDR (CR1), we only observed expression in the context of the N-terminal IDR: we did not observe expression either on its own, or when we replaced the C-terminal IDR in the full length protein. These observations are consistent with the model where the N- and C-termini of Ded1 interact [25], and this is a level of complexity that is beyond our bulk molecular properties paradigm. Deep learning methods can learn to use larger context and higher-order interactions, and have recently revolutionized folded protein design[1,2]. We believe that deep learning can improve intrinsically disordered protein design as well.

## Methods

### A greedy IDR design algorithm to match bulk molecular properties

We first define a bulk molecular property feature space. 94 molecular properties (sequence features, motifs and physicochemical properties) of intrinsically disordered regions were curated from the literature [39,40]. Each property is computed from the primary amino acid sequence using custom python scripts to define a 94-dimensional feature vector for a sequence. Python code to calculate these features is available at https://github.com/IPritisanac/idr.mol.feats [39]

Because the molecular properties include signed integers like net charge, strictly positive fractions like fraction of charged residues, and real numbers like Kyte-Doolittle hydropathy, we scaled the molecular properties by their mean and standard deviation over all the predicted IDRs (obtained from [41]) in the *S. cerevisiae* proteome.

We initialize the designed sequence to a random sequence. Then, at each iteration, for each amino acid in a sequence of length L, we create a sequence with one of the 19 other amino acids, which yields 19*L candidate sequences. We then calculate the Euclidean distance in the (scaled) molecular property feature space between the target sequence and each of the 19*L candidate sequences. We choose the candidate sequence with the smallest distance to target sequence as the designed sequence to start the next iteration. We repeat until the distance to target sequence no longer decreases.

Pseudocode for greedy IDR design:

~~~
target = given by user
L= length(target)
designed_seq = random peptide of length L
Precision = 0.00001
Previous_step_size = 1
Current_distance = calculate distance between designed_seq and target
While Previous_step_size > Precision:
        Candidate_seqs = empty list
        For aa in designed_seq:
                  Change aa to other 19 possibilities and append to candidate_seqs
        all_distances = Calculate 19*L distances to target
        designed_seq= argmin(all_distances)
        Previous_distance = current_distance
        Current distance = min(all_distances)
        Previous_step_size = Current_distance - Previous_distance
Return designed_seq
~~~

### Selection of condensate associated molecular properties

To select features associated with condensate formation for design of the Ded1 N-terminus for condensate formation, we obtained 24 yeast proteins from PhaSepDb [42] and visualized the molecular properties of their IDRs [39]. We found a cluster of these IDRs that showed stereotypical patterns of several molecular properties and we selected 10 of these for further analysis. This approach is expected to identify features common within this select set of condensate-localizing proteins but not features generally associated with all biomolecular condensates.

### Computational analysis of designed IDRs

To confirm that the algorithm was working as expected, for 10 replicates of the S. cerevisiae Cox15 N-terminus design we computed scaled molecular property vectors for each sequence along the design path, starting with the random initialization and ending with final design. To these sequences we added the homologs of the Cox15 N-terminus (obtained from [41]). We performed principal components analysis (PCA, using the R statistics package[43]) and plotted the scores of each sequence on the first two PCs.

We aligned initial 10 random initial peptides, the final designs and Cox15 N-terminus homologs to the S. cerevisiae Cox15 N-terminus (target) sequence using biopython’s pairwise global alignment module[44] with the BLOSUM62 scoring matrix [45] and gap open and extension penalties of 11 and 1, respectively. We computed the percentage of identical residues ignoring gapped positions in the alignment.

MitoFates [29] was accessed at http://mitf.cbrc.jp/MitoFates/cgi-bin/top.cgi The input for the predictor is the full Cox-15 protein sequence. Because the greedy algorithm did not typically produce an M at the start of each sequences, we replaced the Scer Cox-15 residues 2-45 (leaving the initial M) with the corresponding residues from our random initializations and converged peptides designs and submitted them to the MitoFates webserver.

PScore [31] was accessed at http://abragam.med.utoronto.ca/~JFKlab/Software/psp.htm. The input is the full Ded1 protein sequence. We replaced the Ded1 N-terminal IDR segment with each of the random initializations and converged designed peptides and submitted them to the Pscore webserver.

### Yeast strain design

Background strains for both Cox15 and Ded1 serve to show wild-type localization for each protein used in this study. These were made by tagging each ORF with mCherry at their respective endogenous loci via homologous recombination in S. cerevisiae S288C derivative (BY4741: MATa his3Δ1 leu2Δ0 met15Δ0 ura3Δ0) [46] using the standard LiAc method PMID: 17401334. Direct transformation of a linear PCR product containing the codon-optimized mCherry coding sequence and a selectable resistance marker just prior to the native STOP codon yielded C-terminally tagged fusion products which were then isolated by marker selection on yeast synthetic minimal media (SD). These strains both serve as internal control strains, and the full-length ORFs of each remained intact for the duration of the study.

Two-colour experimental IDR strains for both Cox15 and Ded1 - containing full-length sequence or designed IDR sequences - were then made by inserting a second copy of the ORF tagged with SuperFold-GFP (sfGFP)[47] at the HO locus of the corresponding control strain via recombinant homologous transformation as described above. The purpose of this was two-fold, as Ded1 is an essential gene, and substantial alteration of the endogenous gene could inhibit cell growth – and it also provides a consistent internal control for direct comparison of differential localization for designed IDR sequences.

Synthetic amino acid sequences were obtained via computational design and codon optimized for optimal expression in S. cerevisiae (https://www.idtdna.com/CodonOpt)The resultant nucleotide sequences were then used to design gene fragments (gBlocks), (https://www.idtdna.com/site/order/gblockentry), which were used as PCR templates. Linear PCR products were similarly integrated at the HO loci and isolated by marker selection.

Strain genotypes can be found in Table S3.

### Confocal microscopy

Control and mutant strains were inoculated into synthetic minimal media (SD) and grown overnight @ 30°C. Prior to imaging, stationary cultures were diluted 1/10 in fresh media and grown at 30°C for 4 hours to ensure log-phase growth and proper expression of mCherry/GFP fusion products. For Ded1 strains, cultures were split in half, and one half was held at 30°C, while the other half was placed in a water bath at 46°C for 15 minutes. For both Ded1 and Cox15 strains, cells were then fixed in 0.5% paraformaldehyde; 1X PBS pH=7.5. Fixed cells were imaged using a Leica TCS SP8 confocal microscope with a 63X oil-immersion objective and a 10X eye-piece objective. mCherry excitation was at 587nm and GFP excitation was at 488nm. Z-stacks were captured at ∼ 0.2-0.4 μM intervals with pixel size in range of 60-80 nm.

To confirm that paraformaldehyde fixation did not qualitatively affect the condensate localization patterns we observed, we also imaged our Ded1 design strains live after 15 minutes of heat shock and found similar patterns (Supplementary figure S2).

3-D Z-stacks were opened in FIJI[48] and reduced to 2 dimensional images via maximum intensity project for both red and green channels. These images were processed via “despeckle” function in FIJI and saved as RGB TIF images, yielding the mCherry and GFP images. The two channels were then merged in FIJI to produce the “overlay” images.

## Acknowledgements

We thank Drs. Alex Lu, Kevin Yang, Jin Montclare, Michael Elowitz and Audrey Gasch for discussions. We thank Andrew Duncan, Ami Sangster and Drs. Philip Kim, Alex Lu and Taraneh Zarin for comments on the manuscript. This research was supported by project grants from the Canadian Institutes of Health Research (CIHR PJT-148532) and Canada Research Chairs to A.M.M. and J.D.F.-K..

## Supplementary Figures and Tables

**Figure S1.**
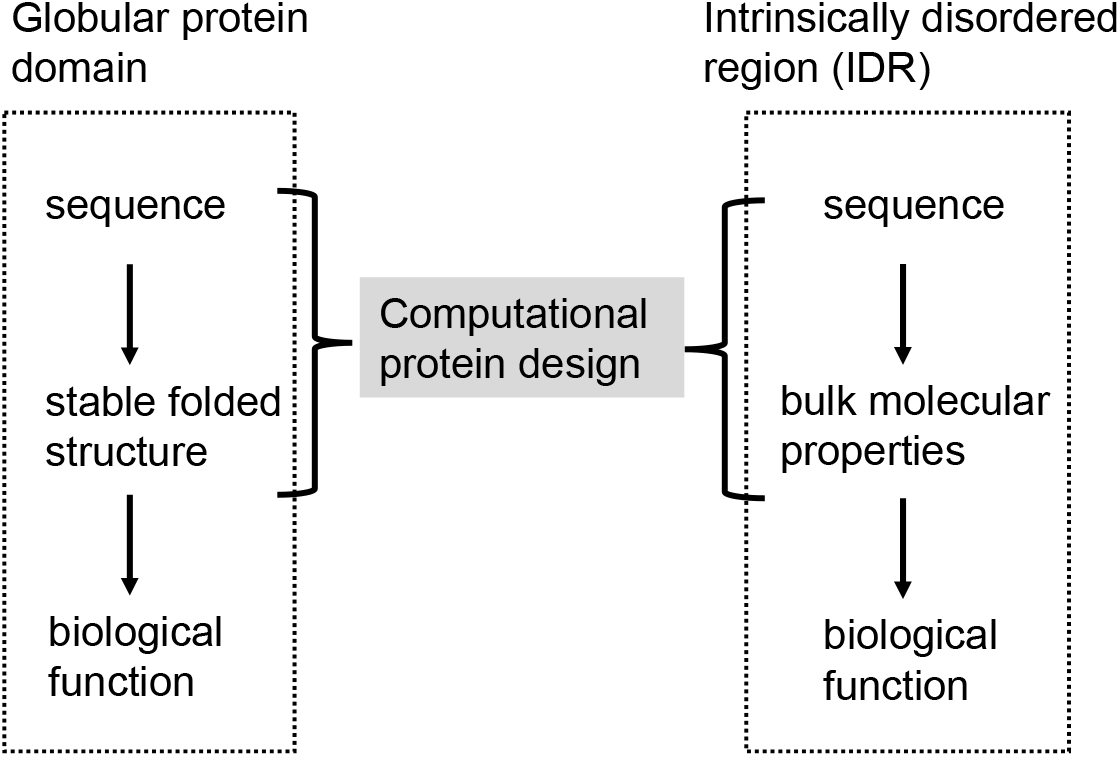
A generic approach to IDR design in analogy to computational design approaches for globular proteins.

**Figure S2.**
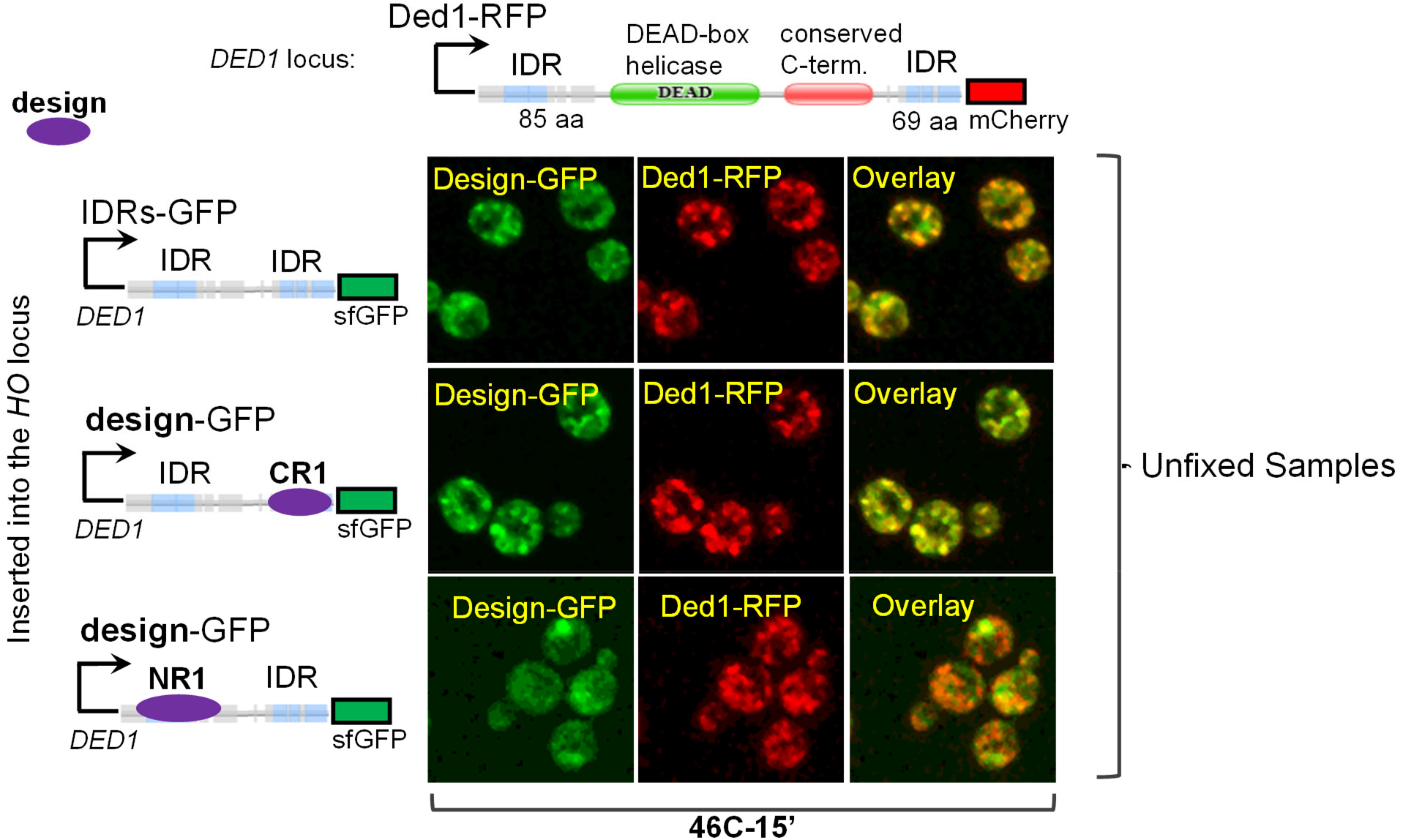
Live cell imaging of Ded1 IDR designs after 15 minutes 46 C heat shock. Strains and images are shown as in Figure 3

**Table S1.**
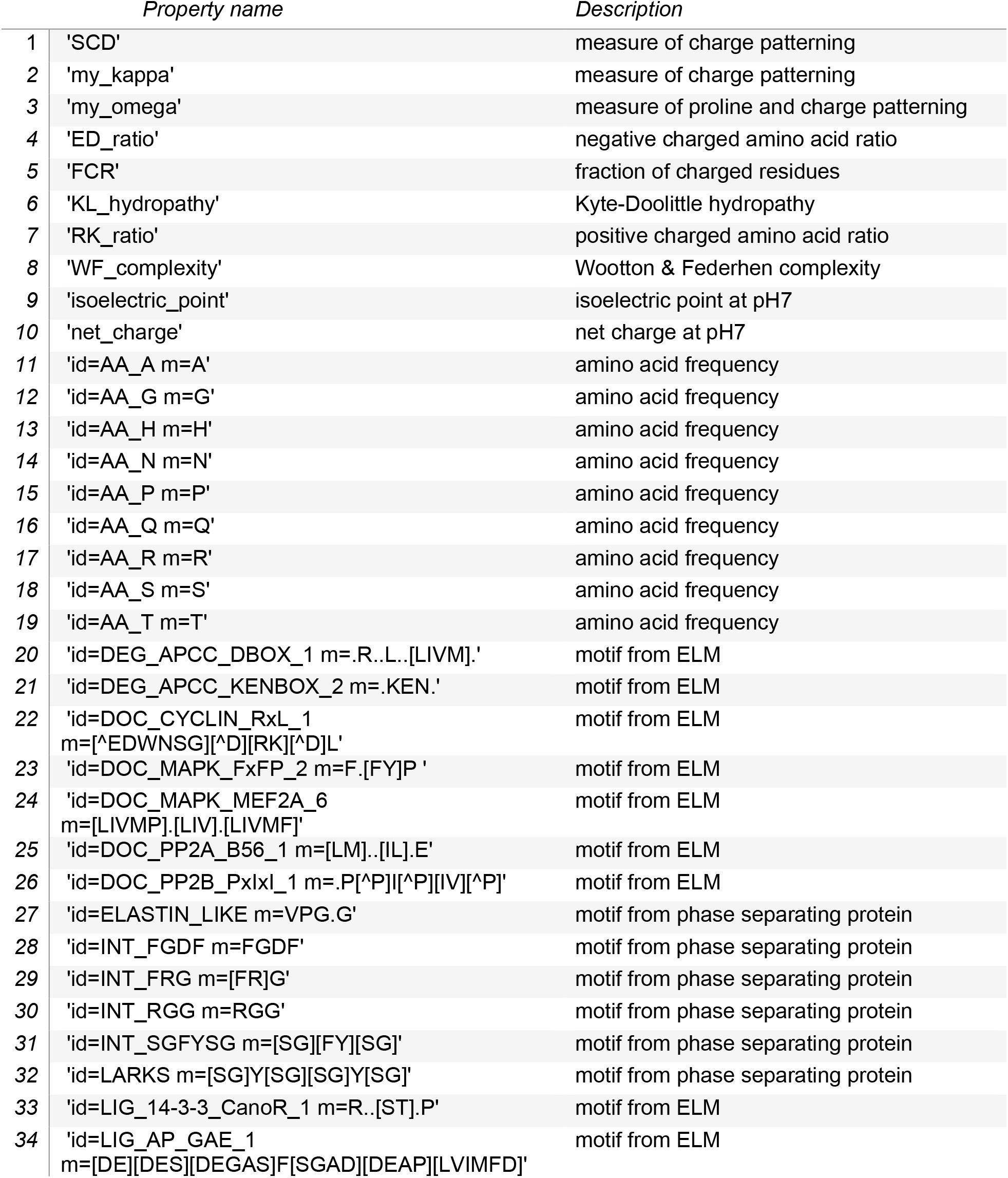
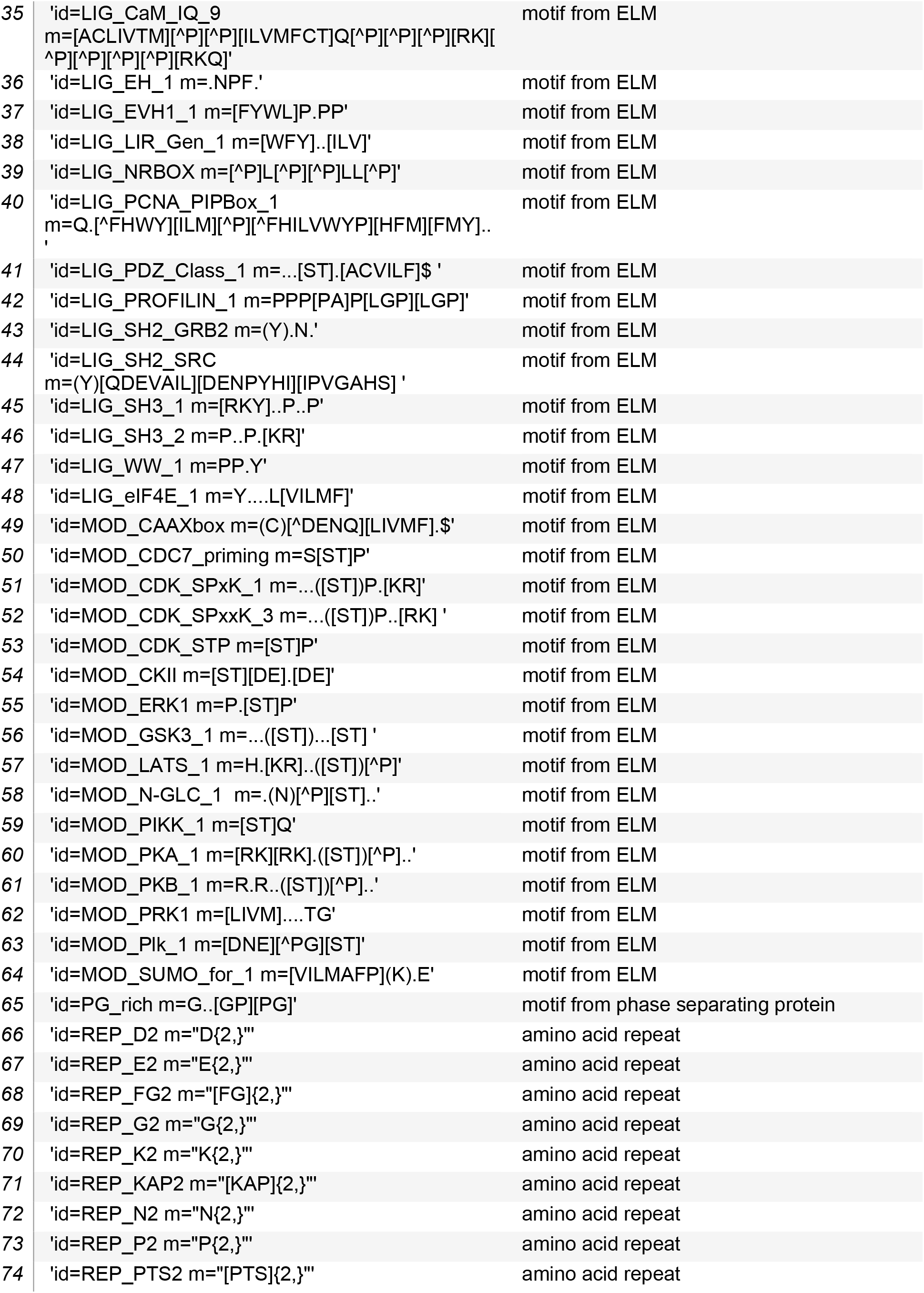
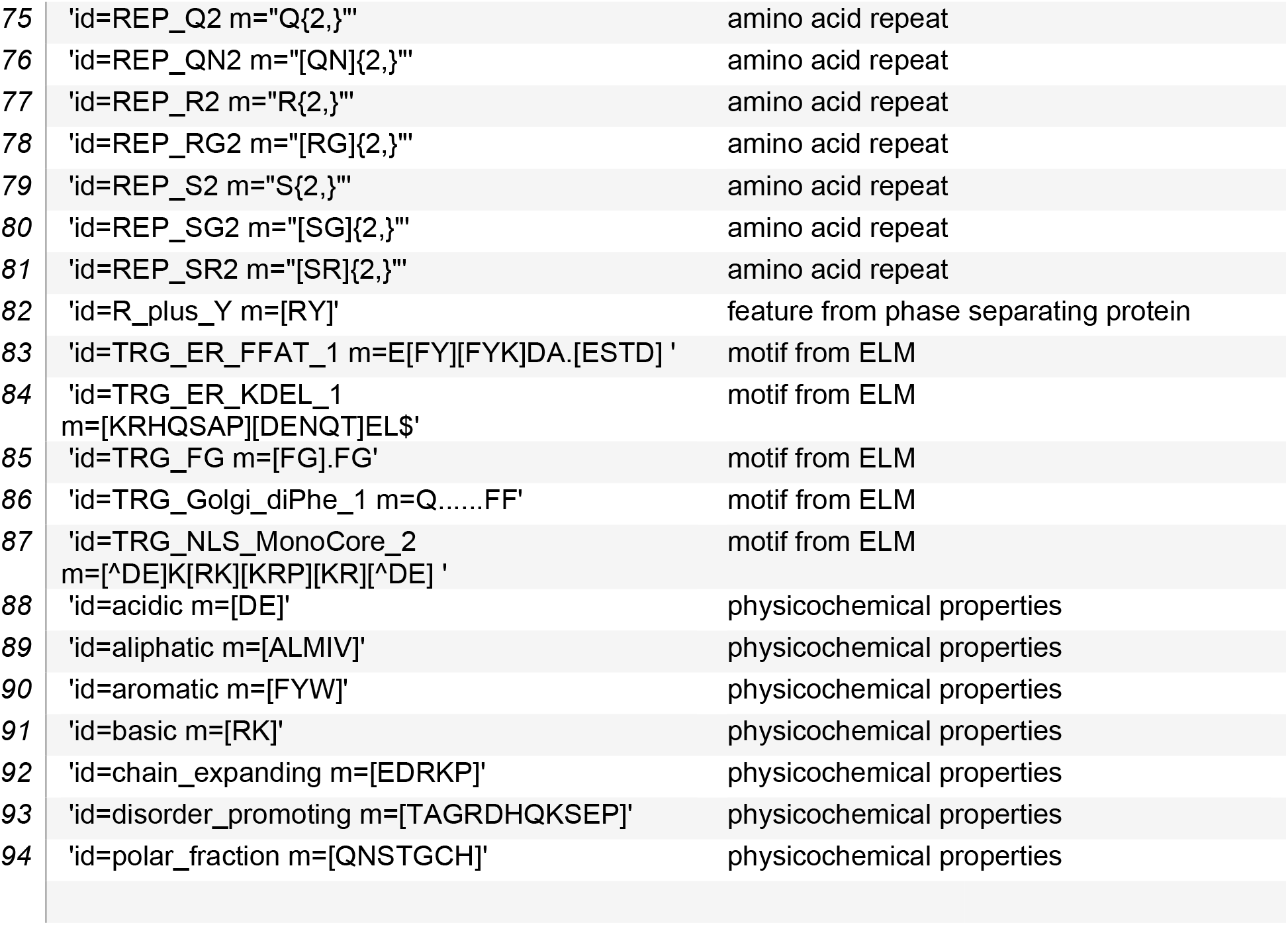
Molecular properties

**Table S2.**
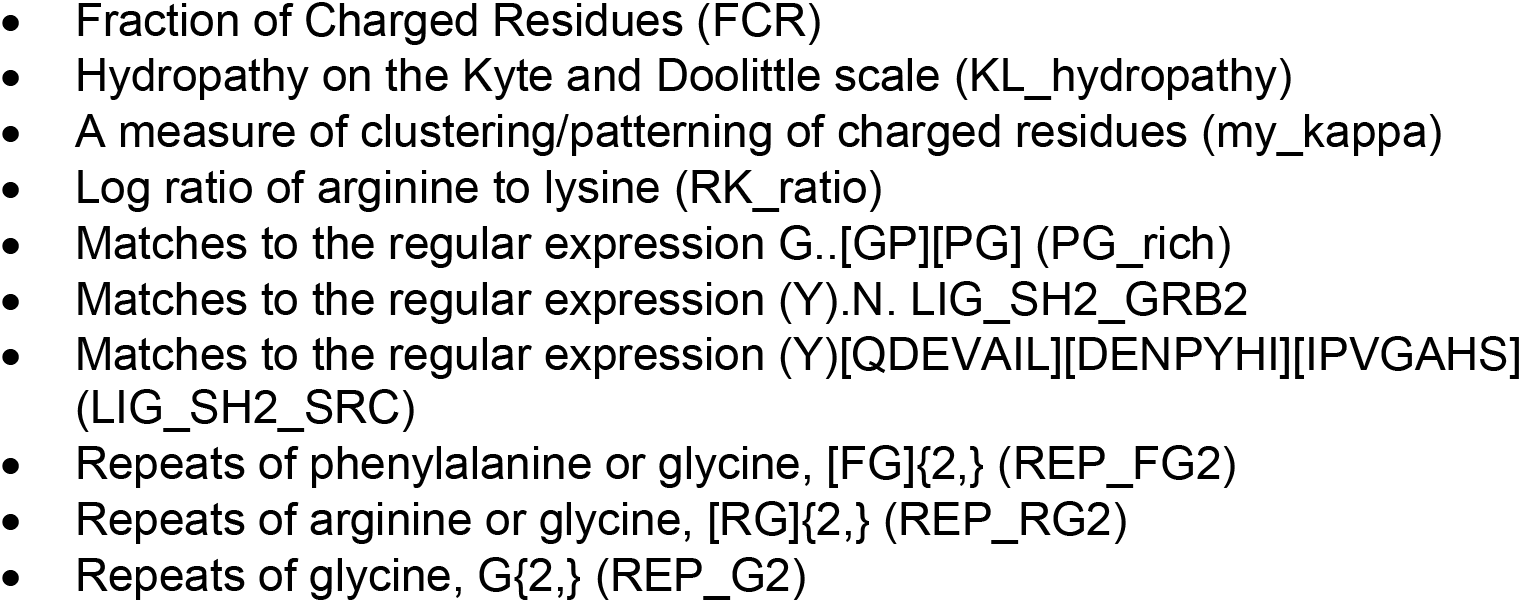
Condensation associated properties derived from a set of condensate-forming proteins (see Methods).

**Table S3.**
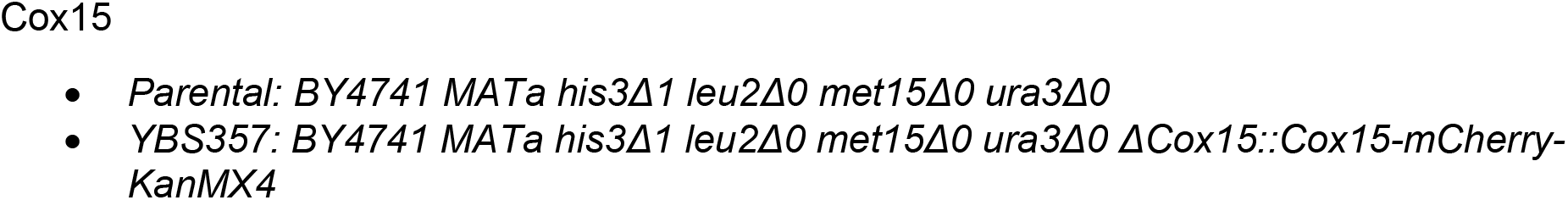
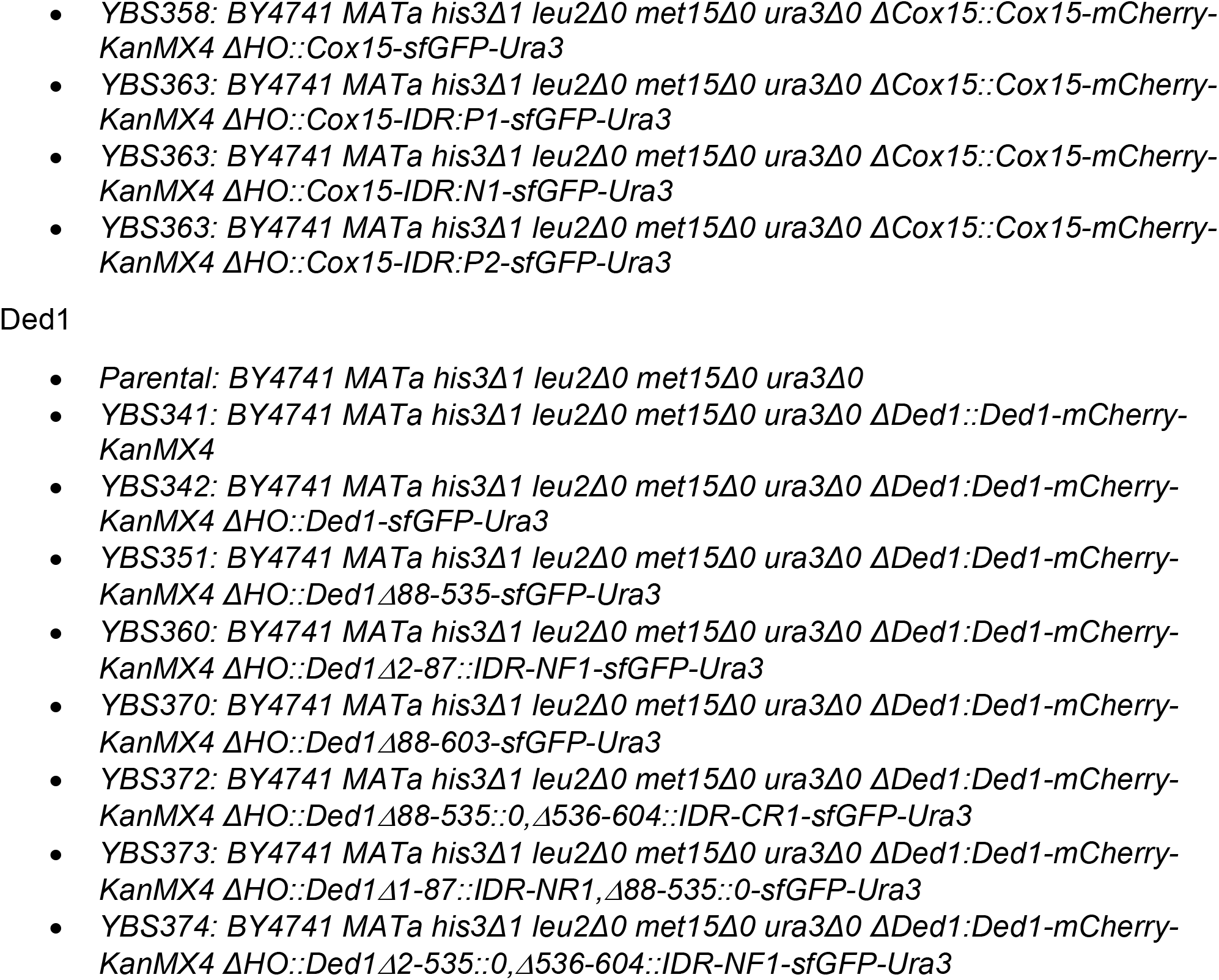
Strain genotypes

